## Supplementary Information for "Accurate and sensitive mutational signature analysis with MuSiCal"

**References** **4**

**2 Supplementary Figures** **5**

### 1 Supplementary Note

#### 1.1 Applicability of mvNMF to tumor somatic mutations

Theoretically, mvNMF is guaranteed to recover the true underlying signatures if the exposure matrix is sparse enough such that only a few signatures are active in each sample, whereas NMF requires much stronger assumptions [1–3]. Indeed, strong sparsity is observed for tumor type-specific exposure matrices produced by PCAWG [4], justifying the application of mvNMF to cancer data (Supplementary Fig. 3a-d). The increased accuracy of mvNMF holds for a wide range of exposure sparsity relevant for cancer data (Supplementary Fig. 3e). Even in the unrealistic case of non-sparse exposures, mvNMF ensures that the solutions are non-negative mixtures of true signatures, enabling the true underlying signatures to be uncovered through the downstream NNLS-based matching step (Supplementary Fig. 4). By contrast, NMF solutions have no such properties and can be arbitrarily distorted. These distortions may result in the discovery of false signatures in the matching step, leading to erroneous signature assignments (Supplementary Fig. 4, also evident from the example in Fig. 2b). Of note, another form of volume regularization has been recently applied to mutational signature discovery for germline mutations [5]. Unlike somatic mutations from tumor samples, the exposure matrix of germline mutations is in general non-sparse due to the lack of heterogeneity, especially when the mutations are analyzed across genomic bins as in [5]. Accordingly, volume regularization is enforced on the exposure matrix in [5]. By contrast, MuSiCal regularizes the volume of the signature matrix (Methods). Therefore, MuSiCal employs the volume regularization that is more suitable for tumor somatic mutations and represents the first application of mvNMF to cancer data.

#### 1.2 Comments on specific signatures related to PCAWG reanalysis

Two ID signatures – ID15 and 16 – in the current COSMIC catalog are not discovered in our PCAWG reanalysis (Extended Data Fig. 7a). ID15 and 16 were found by the PCAWG consortium in three whole-exome sequenced samples from The Cancer Genome Atlas (TCGA) (Extended Data Fig. 7b). However, further inspection shows that ID15 and 16 likely reflect artifactual indels, as these three samples have exceptionally large indel counts ($> 1000$) with only small SBS counts ($< 250$) (Extended Data Fig. 7c). Moreover, their indel variant allele frequencies (VAFs) are much lower than their SBS VAFs, unlike in other TCGA samples (Extended Data Fig. 7d).

COSMIC ID4 is separated into MuSiCal ID4, 19, and 24 in our reanalysis (Extended Data Fig. 7e). Interestingly, our updated MuSiCal version of ID4 is very similar to the experimentally derived signature of topoisomerase 1 (TOP1)-mediated deletions in RNase-H2-null cells, predominantly consisting of 2-bp deletions at short tandem repeats (STRs) with 2 or 3 repeat units, as well as 2-bp deletions with 1-bp microhomology (MH) [6] (Extended Data Fig. 7f). By comparison, the COSMIC version of ID4 contains significant proportions of longer (3- and 4-bp) deletions that are not observed in these experiments, suggesting that COSMIC ID4 is indeed a potential mixture of multiple signatures (Extended Data Fig. 7f).

We analyzed the correlations between per-sample exposures of ID and SBS signatures obtained by MuSiCal and observed strong correlations between ID and SBS signatures with known and related etiologies (Fig. 6c). ID3 correlates with SBS4, both of which are associated with tobacco smoking. The characteristic C deletions at 2- or 3-bp mononucleotide C repeats (i.e., CC or CCC) in ID3 potentially result from repairs of G adducts formed by tobacco carcinogens (Fig. 6a) [7]. ID6 correlates with SBS3, producing $\geq$5-bp deletions with overlapping microhomology (MH), consistent with elevated activities of MH-mediated end joining (MMEJ) in HRD tumors (Fig. 6a, c). ID13, dominated by T deletions at TT dinucleotides, correlates with SBS7 and is associated with UV light exposure (Fig. 6a, c). Several ID signatures, including ID1, 2, and 7, are correlated with MMRD-related SBS14, 15, 20, 21, and 26 (Fig. 6c). ID7 primarily comprises 1- and 2-bp deletions at long repetitive sequences and is known to contribute to a large number of indels in MMRD tumors (Fig. 6a) [4]. ID1 and 2, characterized by 1-bp T insertions and deletions at long homopolymers of T, respectively, reflect higher levels of replication slippage mutagenesis in MMRD tumors (Fig. 6a) [4, 8].

#### 1.3 Impact of over-assignment of SBS40 on other signatures

The impact of the over-assignment of SBS40 cannot be overstated, as it also results in widespread under-assignment of other similar or overlapping signatures. For example, in the results obtained by the PCAWG consortium, SBS8 (unknown etiology) is under-assigned in multiple tumor types where SBS40 is over-assigned (Fig. 7a, Supplementary Fig. 10). Similar under-assignments can be observed for SBS18 (damage by reactive oxygen species), 34 (unknown etiology), and 41 (unknown etiology) (Fig. 7a, Supplementary Fig. 10). To further demonstrate that the over-assignment of SBS40 can confound the interpretation of other signatures, we investigate the correlation between SBS1 and SBS5. Both signatures are referred to as clock-like signatures and correlate with patients' age in most cancers and normal cells [4, 9–11]. We thus expect to observe a correlation between the exposures of SBS1 and 5, although potentially with different slopes in different tumor types. When these tumor type-specific accumulation rates are factored out by multiplying the SBS5 exposure with a normalization factor corresponding to the ratio of average SBS1 and SBS5 exposures for a given tumor type (Supplementary Fig. 13), a constant slope should be observed in the pan-cancer setting. However, signature assignments from the PCAWG consortium show distinct SBS5/SBS1 slopes for samples with and without SBS40 (Fig. 7c, d). In particular, the normalized SBS5/SBS1 ratio is consistently lower in SBS40-positive samples, suggesting that a proportion of mutations caused by SBS5 are misattributed to SBS40 in these samples (Fig. 7c, Supplementary Fig. 13c). By comparison, MuSiCal-derived signature assignments show consistent SBS5/SBS1 slopes that are not confounded by SBS40 (Fig. 7c, d, Supplementary Fig. 13c).

### 2 Supplementary Figures

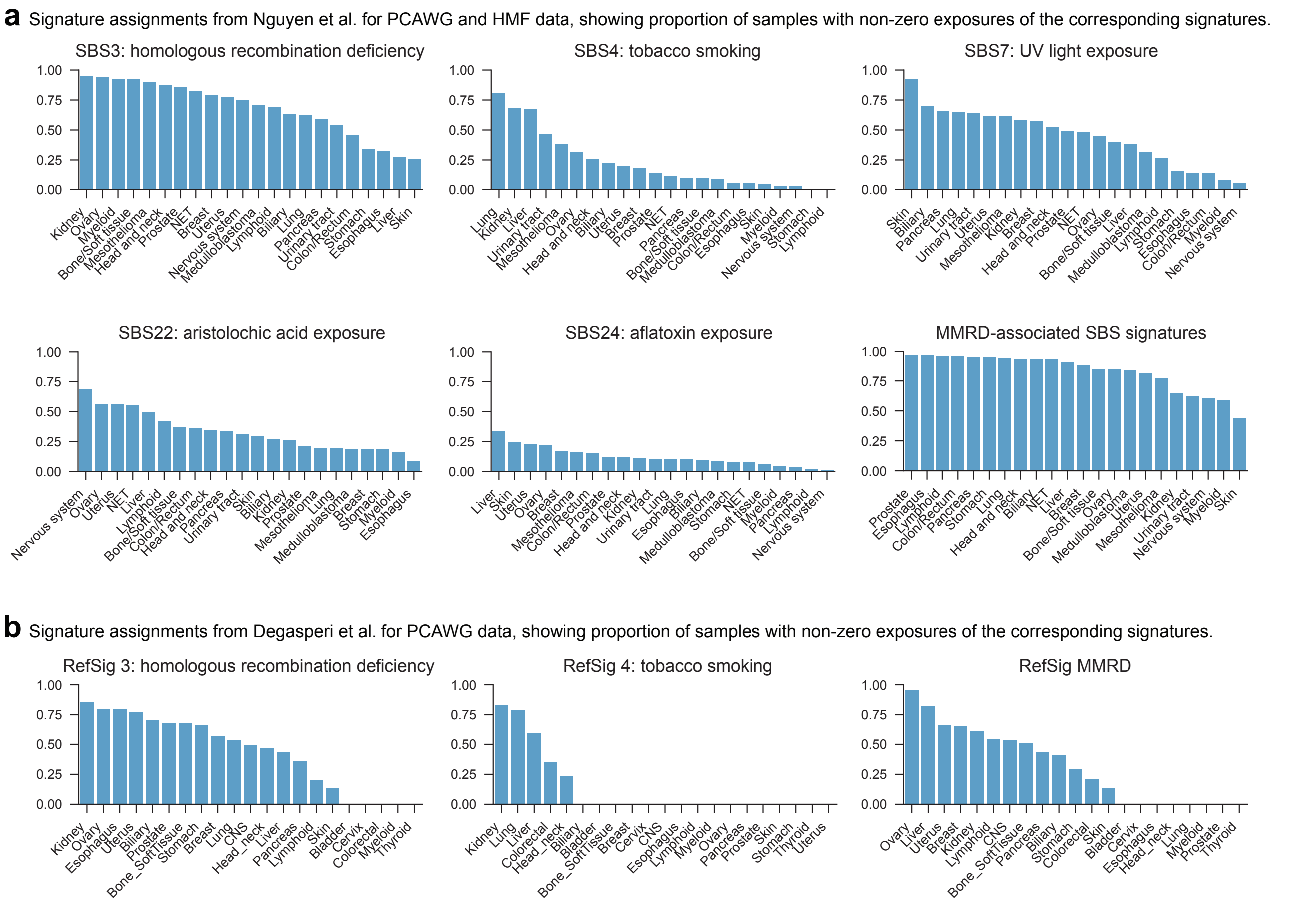

**Supplementary Fig. 1 Examples of erroneous signature assignment in previous studies**. **a.** Examples of signature assignment results obtained from Nguyen et al. [12], showing the proportion of samples with non-zero exposures of the corresponding signatures for each tumor type. Results for PCAWG data and the Hartwig Medical Foundation (HMF) cohort are combined. SBS3 corresponds to homologous recombination deficiency (HRD) but is assigned to a large fraction of samples in many tumor types not known to exhibit HRD, such as kidney, myeloid, etc. Similar problems are observed for other signatures – tobacco smoking-associated SBS4 should be specific to lung, liver, and head and neck cancers; UV light exposure-associated SBS7 should be exclusively observed in skin cancers; aristolochic acid-associated SBS22 and aflatoxin-associated SBS24 should be specific to liver, biliary, and kidney cancers; and MMRD-associated signatures (SBS6, 14, 15, 20, 21, and 26) should be assigned to MMRD samples, which are prevalent only in colorectal, esophagus, stomach, and uterus cancers. **b.** Examples of signature assignment results obtained from Degasperi et al. [13] for PCAWG data. Degasperi et al. use a different catalog of signatures with the prefix RefSig. RefSig 3 is similar to COSMIC SBS3 and corresponds to HRD. RefSig 4 is similar to COSMIC SBS4 and corresponds to tobacco smoking. Two RefSig signatures – RefSig MMR1 and RefSig MMR2 – correspond to MMRD and are combined as RefSig MMRD in the plot. Similar problematic signature assignments as in (a) are observed, e.g., substantial over-assignment of RefSig 3 in many tumor types, RefSig 4 in kidney and colorectal cancers, and RefSig MMRD in tumor types without prevalent MMRD.

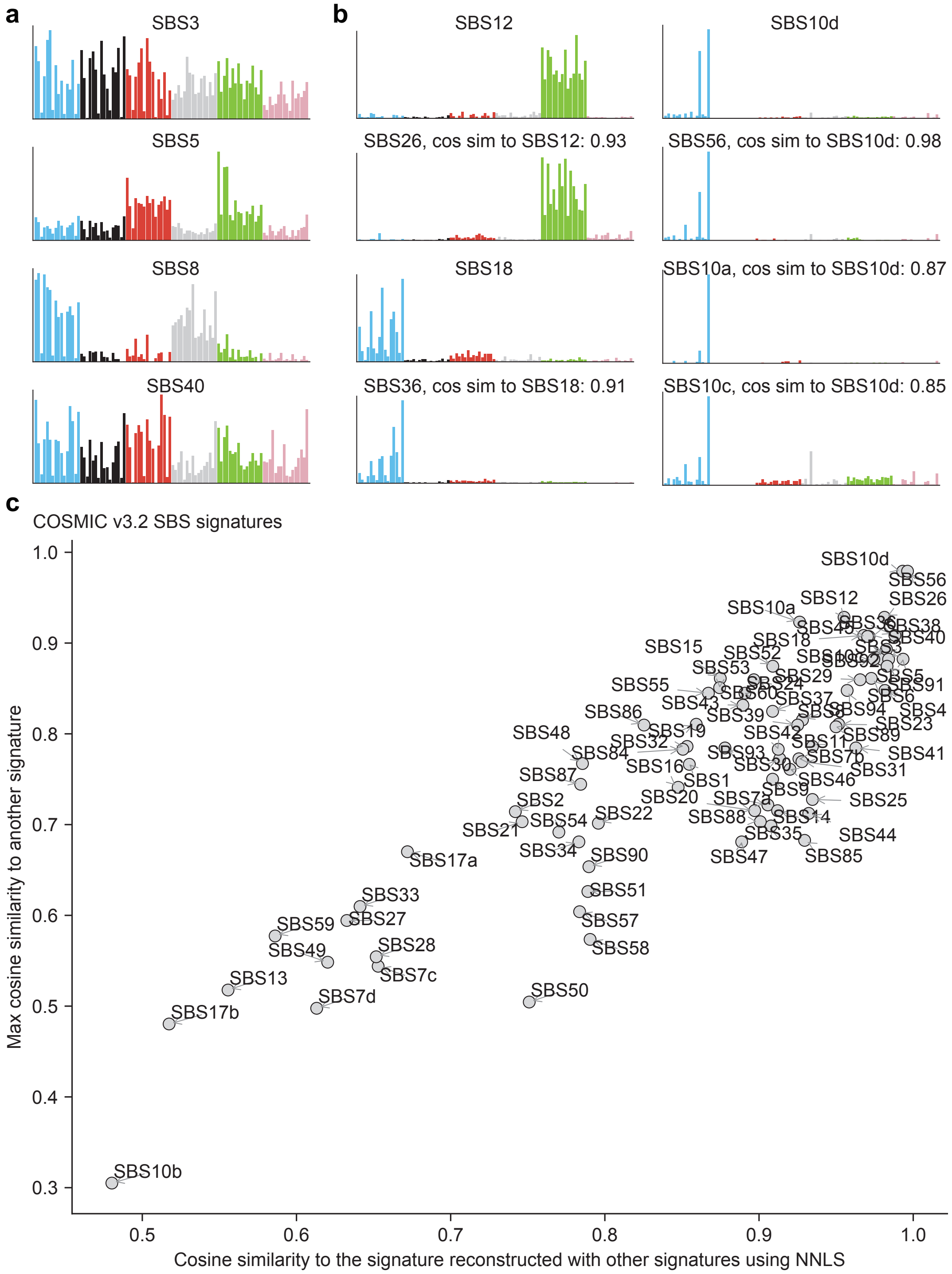

**Supplementary Fig. 2 The COSMIC catalog contains signatures that are flat, overly similar, or can be expressed as linear combinations of each other**. **a.** Examples of relatively flat signatures. **b.** Examples of similar signatures. SBS10d, 56, 10a, and 10c are all similar to each other. SBS12 is similar to SBS26. SBS18 is similar to SBS36. **c.** Similarity between different SBS signatures in the COSMIC catalog. For each signature, the maximum cosine similarity to another signature in the catalog is plotted on the y-axis, and the cosine similarity to the signature reconstructed by fitting with all other signatures in the catalog through NNLS is plotted on the x-axis.

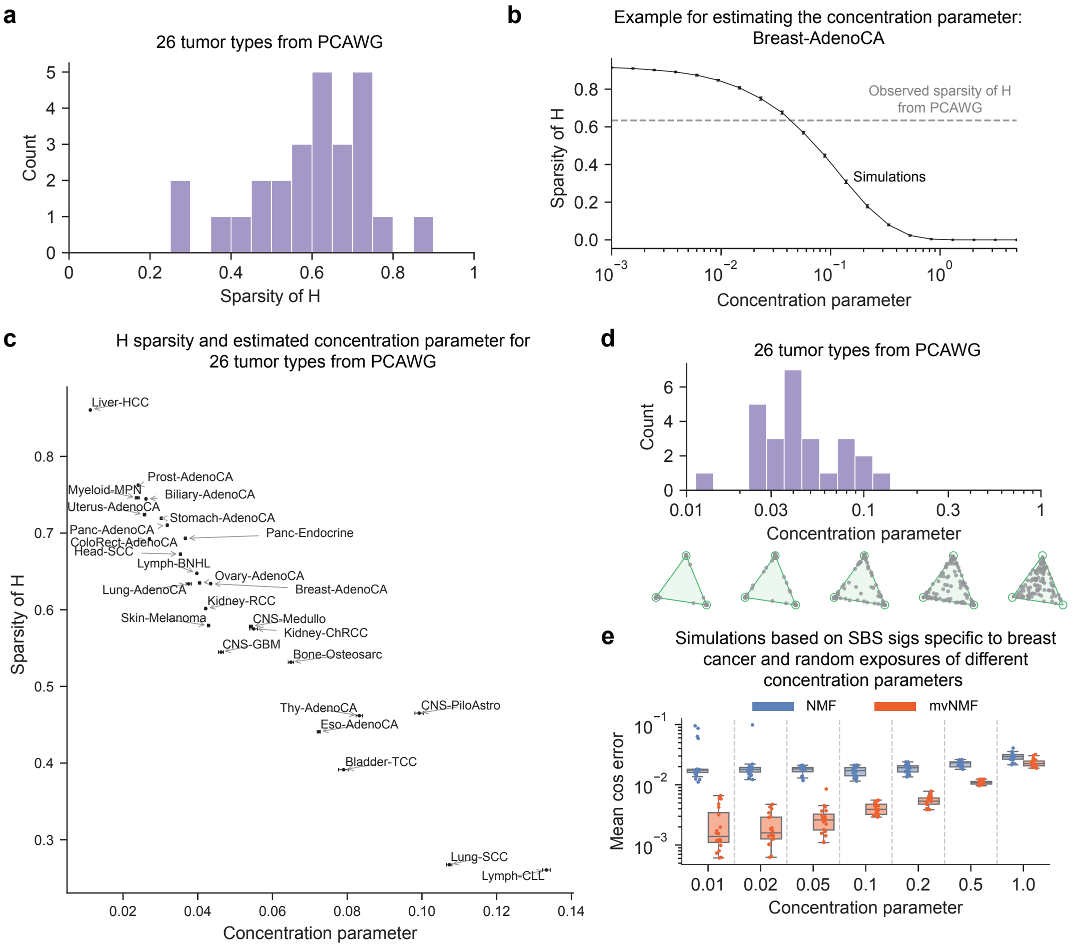

**Supplementary Fig. 3 mvNMF is suitable for *de novo* signature discovery from cancer data.** Theoretically, mvNMF is guaranteed to recover the true underlying signatures if the exposure matrix $H$ satisfies the so-called sufficiently scattered condition [1–3]. Intuitively, the sufficiently scattered condition requires that only a few signatures are active in each sample [1–3]. Although this condition is seemingly reasonable for tumor samples, we still set out to investigate to what extent it is satisfied in real cancer data. Empirically, the sufficiently scattered condition is more likely to be satisfied when $H$ is sparse enough [1, 2]. We therefore study the tumor type-specific exposure matrices from PCAWG results [4]. **a.** Distribution of sparseness (as quantified by the fraction of zero elements) for PCAWG exposure matrices of 26 tumor types with at least 20 samples. The exposure matrices of PCAWG tumors contain 53% of zero elements on average (range 22-86%), indicating strong sparsity. **b.** An example demonstrating how the concentration parameter can be estimated from exposure sparseness. Assuming that the sample exposures in $H$ follow a symmetric Dirichlet distribution, the concentration parameter $\alpha$, defined as the value of the single Dirichlet parameter used in all dimensions, is directly related to the sparseness of $H$. A concentration parameter of 1 results in exposure vectors being uniformly distributed over the simplex, and thus non-sparse, whereas a concentration parameter much smaller than 1 corresponds to sparse exposures. Therefore, in order to obtain a more quantitative insight, we estimate $\alpha$ from exposure sparseness for PCAWG tumors. To illustrate how the estimation is performed, we take the exposure matrix of Breast-AdenoCA from PCAWG results as an example. The exposure matrix has a sparsity level of 0.63 (gray dashed line), as quantified by the fraction of zero elements. Synthetic exposure matrices with the same number of samples and mutation counts are simulated from Dirichlet distributions with different concentration parameters, and the resulting exposure sparseness is plotted (black line, error bar indicating standard deviation of 20 independent simulations). A monotonic and tight relation between exposure sparseness and the concentration parameter is observed, allowing the concentration parameter to be determined using Brent's root-finding algorithm. Because of the randomness from simulations, the solver is run 10 times independently, and the mean solution is taken. For Breast-AdenoCA, $\alpha$ is estimated to be 0.043, with a standard deviation of 0.0002. **c.** Exposure sparseness and estimated concentration parameters (mean values from 10 independent runs) for 26 PCAWG tumor types with at least 20 samples. Error bars indicate the standard deviation of estimated concentration parameters from 10 independent runs. **d.** Distribution of estimated concentration parameters for PCAWG exposure matrices of 26 tumor types with at least 20 samples. Low-dimensional examples are also shown to illustrate the sparsity level at different concentration parameters of 0.01, 0.03, 0.1, 0.3, and 1, respectively. The estimated $\alpha$ is 0.05 on average (range 0.01-0.13) for PCAWG exposure matrices – much smaller than 1 – again indicating strong sparsity. Therefore, somatic mutations from cancer data are highly likely to satisfy the sufficiently scattered condition, justifying the application of mvNMF. **e.** mvNMF improves the accuracy of *de novo* signature discovery for a wide range of exposure sparseness relevant for cancer data. Simulations in Fig. 2c and Extended Data Fig. 1 are repeated for different $\alpha$ values between 0.01 and 1. An example is shown for simulations based on SBS signatures specifically present in breast cancer. Cosine errors averaged across these signatures are plotted for different values of concentration parameters.

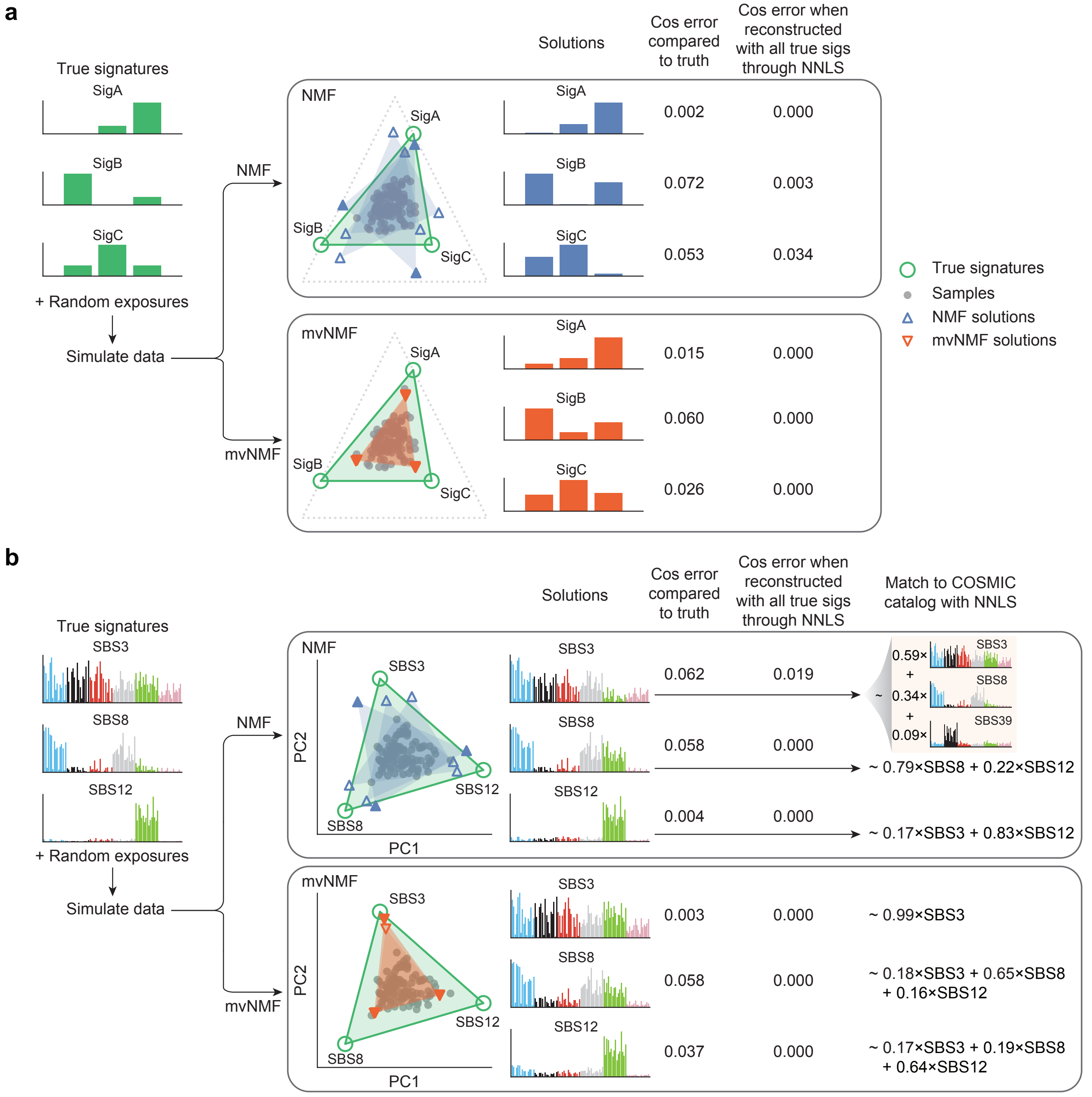

**Supplementary Fig. 4 mvNMF is favored over NMF even when the exposure matrix is non-sparse, as mvNMF always ensures that the solutions are non-negative mixtures of true signatures. a.** An example with three-dimensional signatures. Synthetic samples are simulated as in Fig. 2a, except that a large concentration parameter $\alpha = 5$ is used for Dirichlet-distributed random exposures. NMF and mvNMF are then applied to recover the signatures with three different initializations (shared between NMF and mvNMF). The resulting signature solutions are visualized on the simplex $x + y + z = 1$ as scatter plots. The spectra of example solutions (highlighted by filled triangles for NMF and mvNMF solutions) are shown. In this example, the exposure matrix is dense enough such that all signatures are active in each single sample. As a result, the signatures discovered by mvNMF can be inaccurate if directly compared to the underlying truth due to shrinkage from the volume regularization (see the cosine errors between the solutions and the corresponding true signatures annotated next to the example spectra). However, the mvNMF solutions are guaranteed to be non-negative mixtures of the true signatures, as indicated by the zero cosine errors between the solutions and the signatures reconstructed with all three true signatures through NNLS. By contrast, NMF solutions have no such guarantees: they can be arbitrarily distorted and thus cannot be reconstructed with the true signatures through NNLS. **b.** An example with real SBS signatures. The simulations are performed as in (a), except that real SBS signatures (SBS3, 8, and 12) are used, and the signature solutions are visualized in the PCA space as scatter plots. Again, mvNMF solutions are ensured to be non-negative mixtures of true signatures, whereas NMF solutions are not. Consequently, when matched to the entire COSMIC catalog through NNLS (with an exposure cutoff of 0.05), mvNMF solutions are correctly identified as combinations of true signatures. By contrast, an additional false signature SBS39 is identified for the NMF solution of SBS3 to compensate for the negative algorithmic distortions.

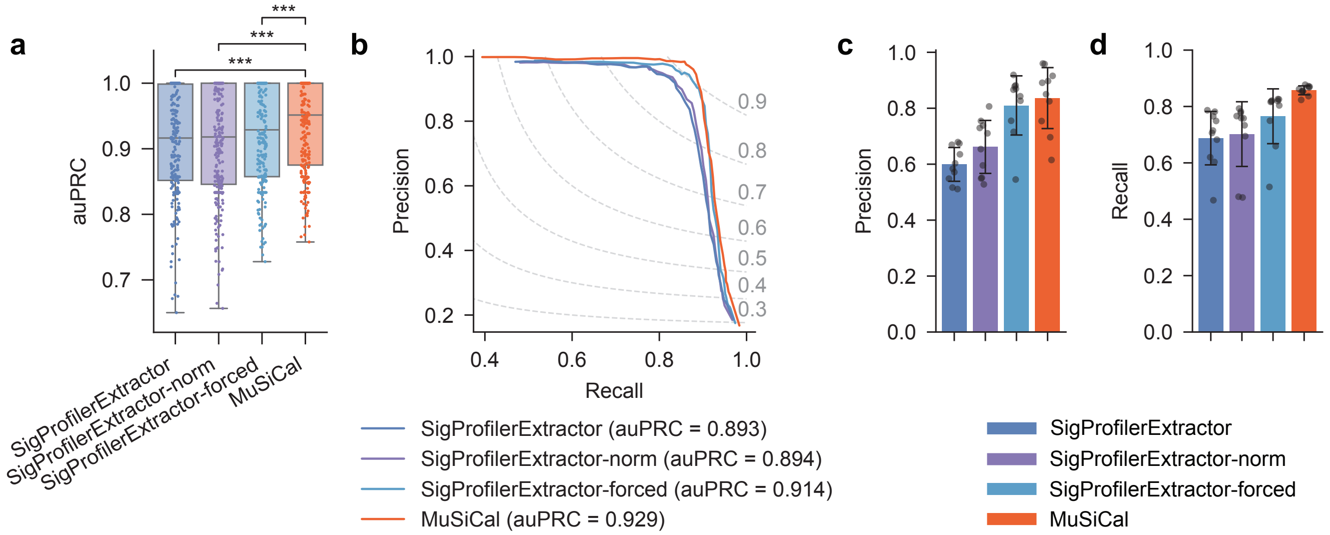

**Supplementary Fig. 5 MuSiCal outperforms the state-of-the-art algorithm SigProfilerExtractor for *de novo* signature discovery.** Here we further benchmark the performance of MuSiCal for *de novo* signature discovery against two variations of SigProfilerExtractor. In one variation, SigProfilerExtractor's built-in input normalization is turned on (`-nx gmm`), and the corresponding result is denoted as SigProfilerExtractor-norm. In the other variation, SigProfilerExtractor is forced to select the same number of signatures as MuSiCal, and the corresponding result is denoted as SigProfilerExtractor-forced. **a.** Area under precision-recall curve (auPRC) for MuSiCal, SigProfilerExtractor, SigProfilerExtractor-norm, and SigProfilerExtractor-forced. Each box in the box plot represents 250 synthetic datasets (25 tumor types $\times$ 10 replicates). auPRC was calculated for each dataset separately, as in Fig. 3b. ***: $p < 0.0005$. $p$-values were calculated with two-sided paired t-tests. Raw $p$-values from top to bottom: $2.8 \times 10^{-4}$, $2.7 \times 10^{-9}$, $7.5 \times 10^{-10}$. **b.** Precision-recall curve (PRC) for the four algorithms. Each PRC represents the average result of 250 synthetic datasets (25 tumor types $\times$ 10 replicates), as in Fig. 3c. **c.** Precision of the four algorithms averaged across all tumor types. Recall was fixed at 0.9. Error bars indicate standard deviation over 10 replicates. **d.** Recall of the four algorithms averaged across all tumor types. Precision was fixed at 0.98, corresponding to a false discovery rate (FDR) of 2%. Error bars indicate standard deviation over 10 replicates.

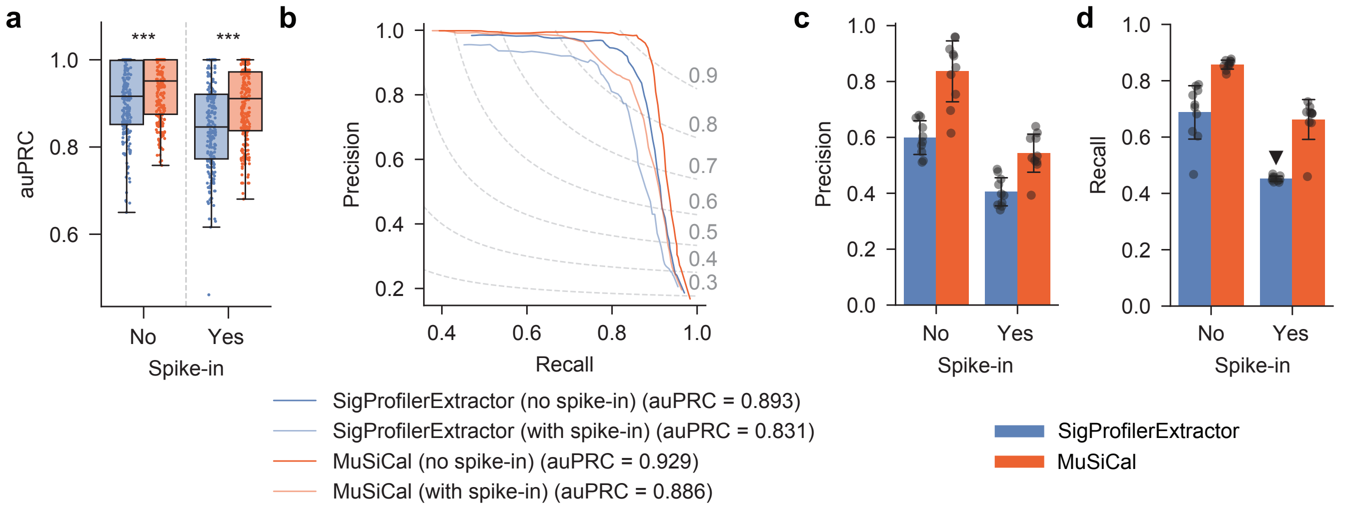

**Supplementary Fig. 6 MuSiCal outperforms SigProfilerExtractor for *de novo* signature discovery when an unknown spurious signature is present.** In order to further investigate the robustness of *de novo* signature discovery by MuSiCal and SigProfilerExtractor, additional mutations from COSMIC SBS48 were spiked-in to the synthetic datasets used in Fig. 3 and Supplementary Fig. 5. The number of spike-in mutations were equal to 5% of the mutational burden for each sample. SBS48 was chosen to represent a spurious signature because it is not present in the original synthetic dataset and is annotated in COSMIC as a possible sequencing artifact found in cancer samples that were subsequently blacklisted for poor quality of sequencing data. To pretend that SBS48 was unknown, we removed SBS48 from the COSMIC catalog when matching the *de novo* signatures and calculating precision and recall. **a.** Area under precision-recall curve (auPRC) for MuSiCal and SigProfilerExtractor with and without spike-in. Each box in the box plot represents 250 synthetic datasets (25 tumor types $\times$ 10 replicates). auPRC was calculated for each dataset separately, as in Fig. 3b. ***: $p < 0.0005$. $p$-values were calculated with two-sided paired t-tests. Raw $p$-values from left to right: $7.5 \times 10^{-10}$, $1.8 \times 10^{-15}$. **b.** Precision-recall curve (PRC) for MuSiCal and SigProfilerExtractor with and without spike-in. Each PRC represents the average result of 250 synthetic datasets (25 tumor types $\times$ 10 replicates), as in Fig. 3c. **c.** Precision of MuSiCal and SigProfilerExtractor averaged across all tumor types with and without spike-in. Recall was fixed at 0.9. Error bars indicate standard deviation over 10 replicates. **d.** Recall of MuSiCal and SigProfilerExtractor averaged across all tumor types with and without spike-in. Precision was fixed at 0.98, corresponding to a false discovery rate (FDR) of 2%. The black triangle indicates the case where a precision of 0.98 was never achieved and the recall at the highest achieved precision was shown. Error bars indicate standard deviation over 10 replicates.

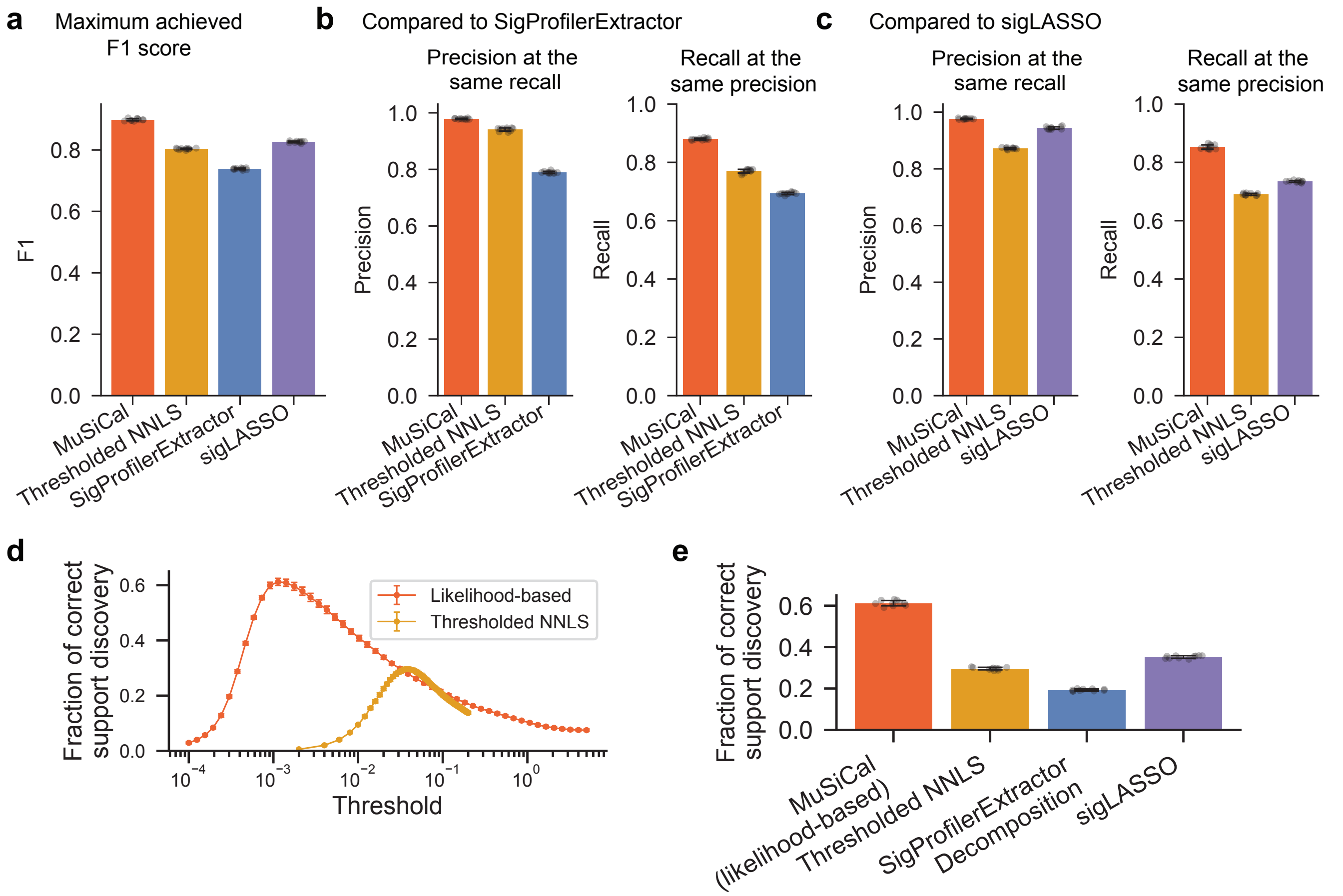

**Supplementary Fig. 7 MuSiCal (with likelihood-based sparse NNLS) outperforms thresholded NNLS, SigProfilerExtractor Decomposition, and sigLASSO for refitting.** The same benchmark results as in Fig. 4c are used. **a.** The largest achieved F1 score of MuSiCal and thresholded NNLS in comparison to the F1 score of SigProfilerExtractor Decomposition and sigLASSO achieved with default parameters. **b.** Precision of MuSiCal and thresholded NNLS at the same recall as SigProfilerExtractor Decomposition (left) and recall of MuSiCal and thresholded NNLS at the same precision as SigProfilerExtractor Decomposition (right). **c.** Precision of MuSiCal and thresholded NNLS at the same recall as sigLASSO (left) and recall of MuSiCal and thresholded NNLS at the same precision as sigLASSO (right). **d.** Instead of precision and recall, an alternative metric of correct support discovery rate – defined as the proportion of samples that have the correct set of active signatures identified – can be used. For this, the thresholds of likelihood-based sparse NNLS and thresholded NNLS are tuned such that the best correct support discovery rate is achieved. **e.** Correct support discovery rate of all four algorithms. Results from MuSiCal (i.e., likelihood-based sparse NNLS) and thresholded NNLS are shown with optimal thresholds. In all panels, data are represented as mean values ± standard deviations (error bars) from 10 independent simulations.

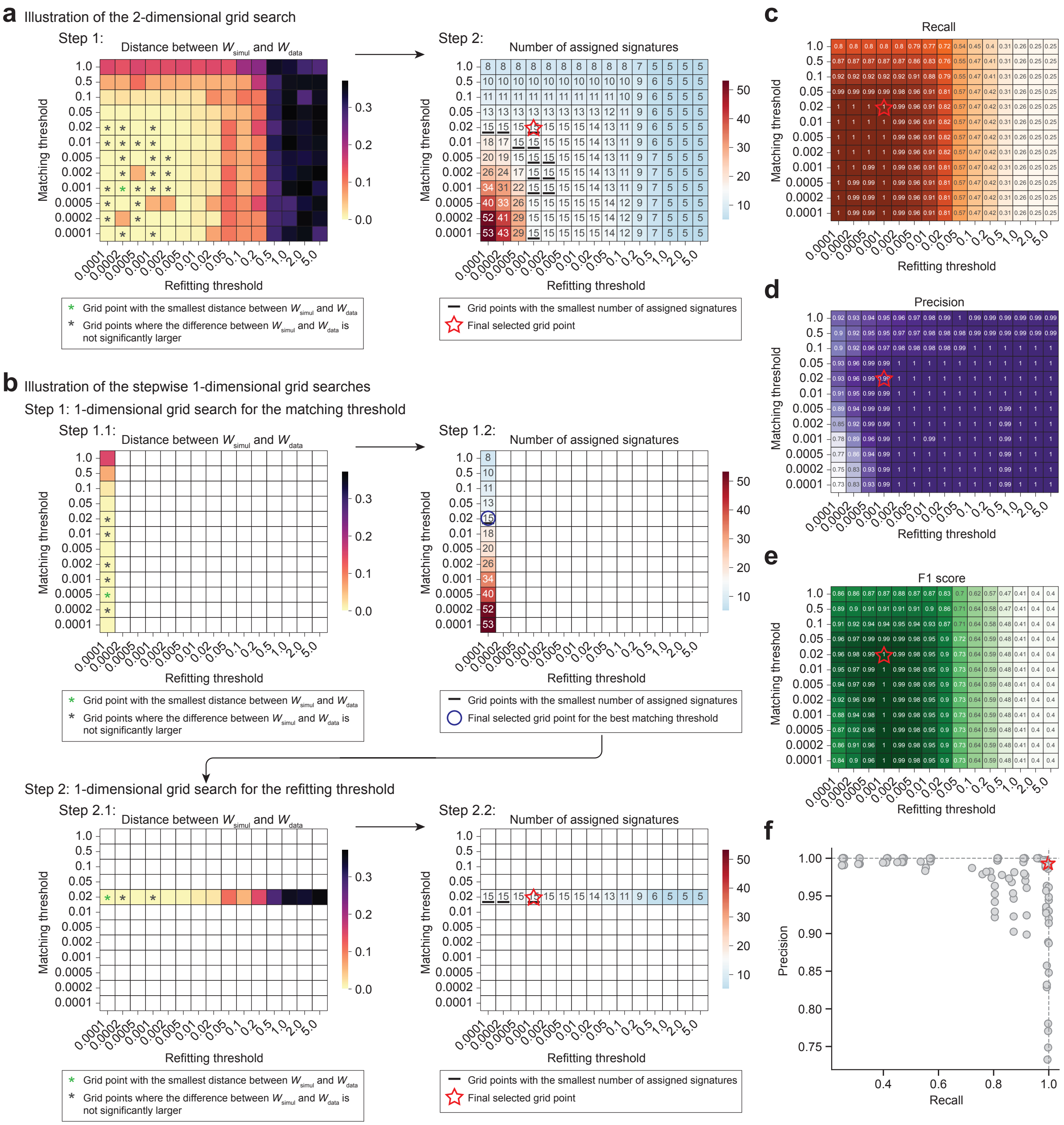

**Supplementary Fig. 8** (Continued on the following page.)

**Supplementary Fig. 8 An example based on the PCAWG Skin Melanoma dataset to illustrate post-discovery parameter optimization through data-driven simulations.** The *in silico* validation/optimization module can be used to select the optimal post-discovery parameters by comparing the original data with data-driven simulations. To demonstrate this approach, a realistic synthetic dataset based on PCAWG skin melanomas is used, such that the true underlying signatures (15 signatures) and exposures are known to evaluate the results. An initial *de novo* signature discovery is first performed to derive $W_{\text{data}}$ and $H_{\text{data}}$ (see main text and Fig. 5a for explanation of the notations). Then, the two likelihood thresholds used in matching and refitting, respectively, are optimized through a two-dimensional grid search with data-driven simulations. Specifically, for each grid point (i.e., pair of likelihood thresholds), $X_{\text{simul}}$ is simulated from the corresponding signature assignments $W_{\text{s}}$ and $H_{\text{s}}$ and then subject to *de novo* signature discovery again, resulting in $W_{\text{simul}}$ and $H_{\text{simul}}$. The optimal grid point is selected based on the comparison between $W_{\text{simul}}$ and $W_{\text{data}}$. Alternatively, to reduce computation time, two separate one-dimensional grid searches can be performed for the matching and refitting thresholds, respectively. **a.** Illustration of the two-dimensional grid search for jointly optimizing matching and refitting thresholds. In the first step, we inspect the mean cosine distance between $W_{\text{simul}}$ and $W_{\text{data}}$ (left panel). The grid point with the smallest distance (marked by green '*') is selected as a candidate solution. Additional grid points (marked by black '*') are further included in the candidate list if the corresponding difference between $W_{\text{simul}}$ and $W_{\text{data}}$ is not significantly larger (see Methods for details). In the second step, we inspect the number of assigned signatures (right panel). Among the candidate grid points selected in the first step, the one with the smallest number of assigned signatures is selected. In this case, there are multiple such grid points (marked by '−'). We select the one with the largest thresholds, i.e., the sparsest signature assignment (marked by a red star). **b.** Illustration of the alternative approach using stepwise one-dimensional grid searches. In the first grid search (top two panels), we optimize the matching threshold while fixing the refitting threshold to a small value (0.0001). In the second grid search (bottom two panels), we further optimize the refitting threshold while fixing the matching threshold to its optimal value (0.02). The best grid point is selected with the same procedure described in (a) for both grid searches. In this example, the joint approach (a) and the stepwise approach (b) select the same optimal grid point. **c-e.** Recall (c), precision (d), and F1 score (e) of signature assignments at different grid points. Recall and precision are calculated by comparing nonzero or zero entries in $H_{\text{s}}$ with those in the true exposure matrix. The selected optimal grid point in (a) and (b) maximizes the F1 score, justifying the approach of parameter optimization by comparing $W_{\text{simul}}$ and $W_{\text{data}}$. Note that different pairs of likelihood thresholds for matching and refitting can lead to the same signature assignment results. Therefore, multiple grid points achieve the best F1 score. **f.** Precision vs. recall of signature assignments at different grid points.

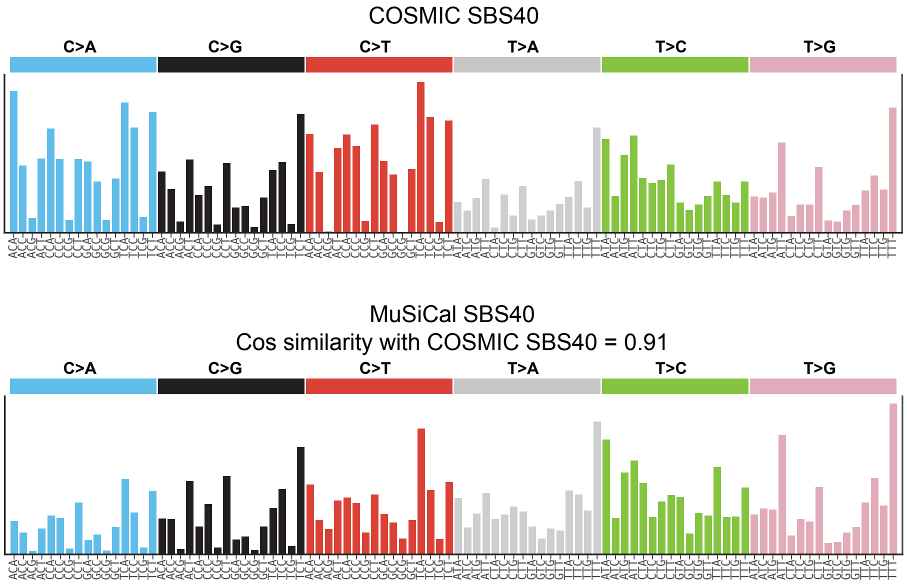

**Supplementary Fig. 9  MuSiCal discovers a modified version of SBS40.** SBS40 in the COSMIC catalog (top) is compared to the SBS40 discovered from our PCAWG reanalysis with MuSiCal (bottom). Compared to COSMIC SBS40, MuSiCal SBS40 has higher proportions of T>N mutations.

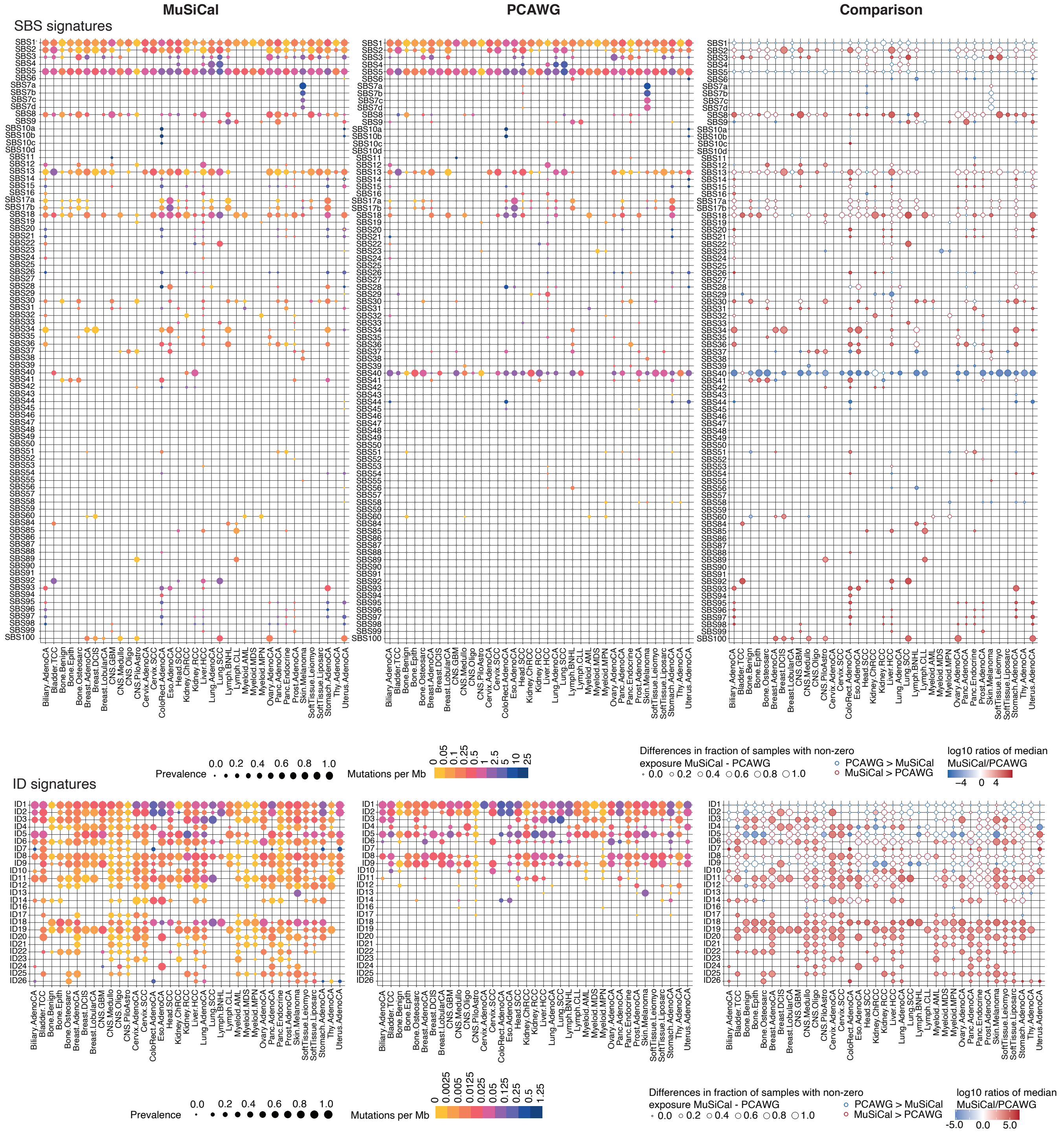

**Supplementary Fig. 10 Comparison of signature assignments obtained by MuSiCal and the PCAWG consortium.** Left panel is the same as Fig. 6b and included for visual comparison. Middle panel is obtained with exposure matrices produced by the PCAWG consortium [4], except that empty rows are added for the new signatures discovered by MuSiCal (SBS95-100 and ID19-27) as well as signatures recently added to the COSMIC catalog since the publication of [4] (SBS84-94). Note that exposures of ID11a and 11b are added together and presented as ID11 for MuSiCal results in order to facilitate comparison with PCAWG results. Right panel shows the comparison of MuSiCal and PCAWG assignments, where marker size represents the difference in proportion of samples with nonzero exposures of the corresponding signature for a given tumor type, and color indicates the log10 ratio of the median number of mutations per Mb contributed by the corresponding signature among samples with nonzero exposures.

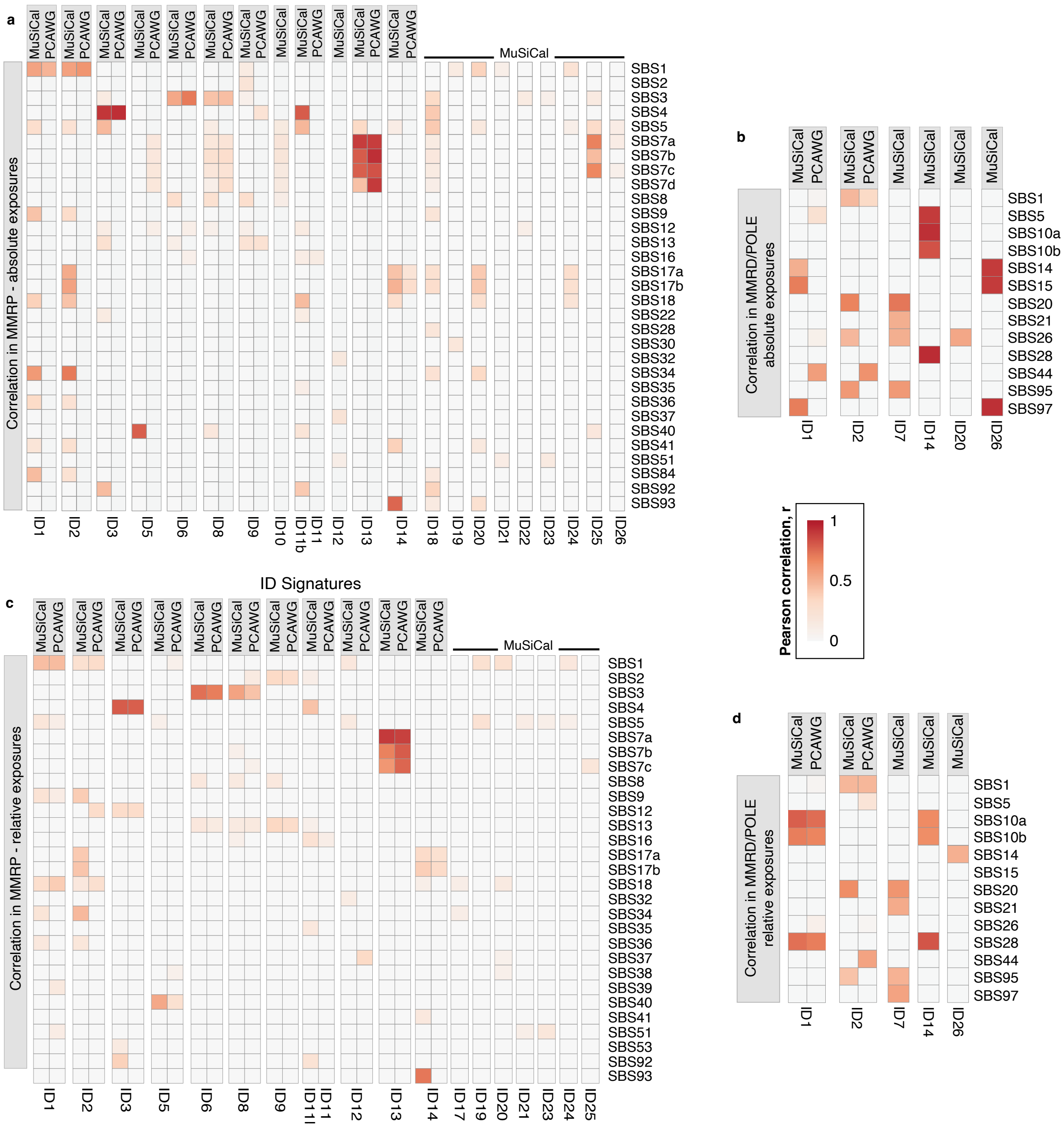

**Supplementary Fig. 11 Comparison of SBS-ID associations in MuSiCal- and PCAWG-derived signature assignments.** **a.** Same as Fig. 6c, except that correlations obtained using signature assignments produced by the PCAWG consortium are also shown for comparison for MMRP samples. **b.** Same as panel (a) but for MMRD, MMRD + POLE-exo mutant or POLE-exo mutant samples. **c-d.** Same as panels a-b, respectively, but for relative ID and SBS signature exposures.

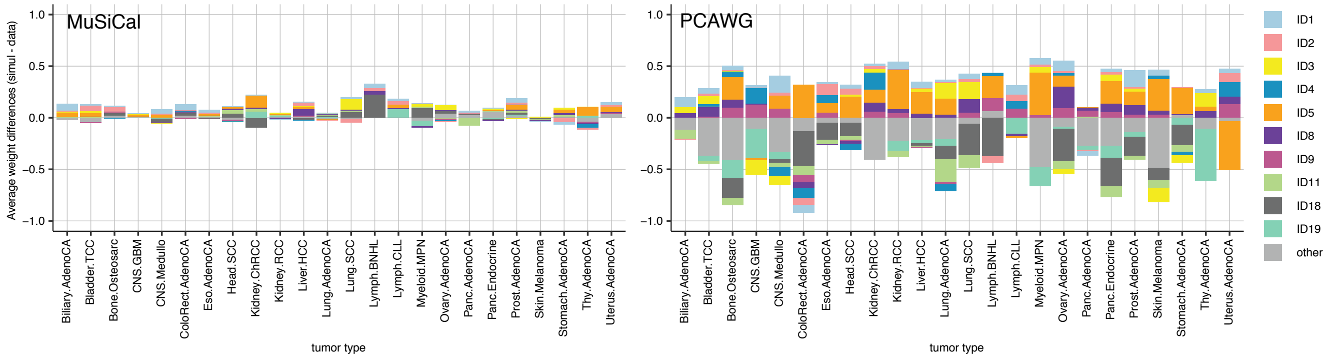

**Supplementary Fig. 12 Comparison between MuSiCal- and PCAWG-derived ID signature assignments in terms of their consistency with data.** Same as Fig. 7a, but for ID signatures.

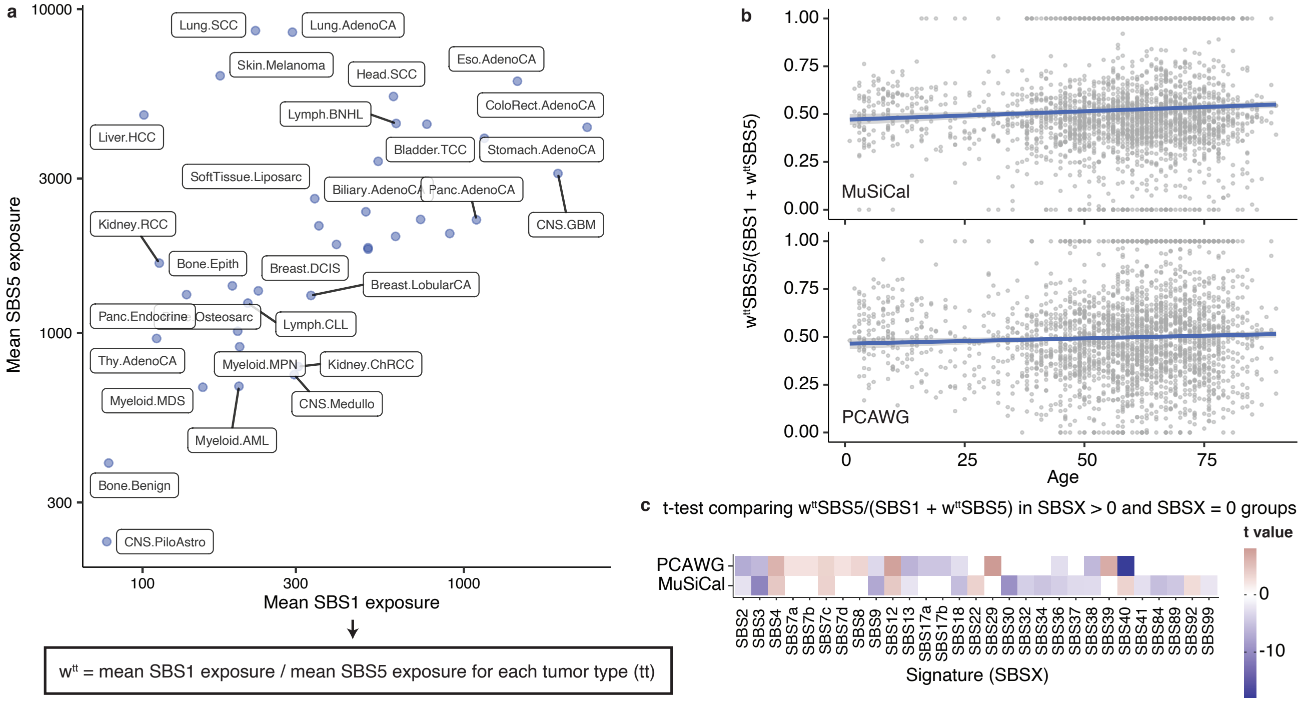

**Supplementary Fig. 13 Correlation between clock-like signatures, SBS1 and 5. a.** Mean SBS5 exposure vs. mean SBS1 exposure in different tumor types, as obtained by MuSiCal, demonstrating the tumor type-specific accumulation rates of mutations from SBS1 and 5. To account for this tumor type-specificity, a normalization factor $w^{tt}$ is calculated as the ratio between mean SBS1 exposure and mean SBS5 exposure for each tumor type ($tt$). The per-sample exposure of SBS5 is multiplied by $w^{tt}$ from the corresponding tumor type when compared with that of SBS1. Exposures obtained by the PCAWG consortium are normalized in a similar way. **b.** Fraction of normalized SBS5 exposure relative to the sum of SBS1 and normalized SBS5 exposures vs. patient age. Since both SBS1 and SBS5 are clock-like and correlated, this fraction is expected to be a constant with respect to age and have a tight distribution (Fig. 7d). Error bands (shaded areas) indicate 95% confidence intervals of the linear regressions. **c.** Comparison of samples with non-zero and zero exposures of different signatures suggests that SBS40 is the major confounder in the signature assignments obtained by the PCAWG consortium. Two-sided t-tests are performed to compare the fraction shown in (b) between samples with non-zero and zero exposures of a signature. Only significant results are shown.

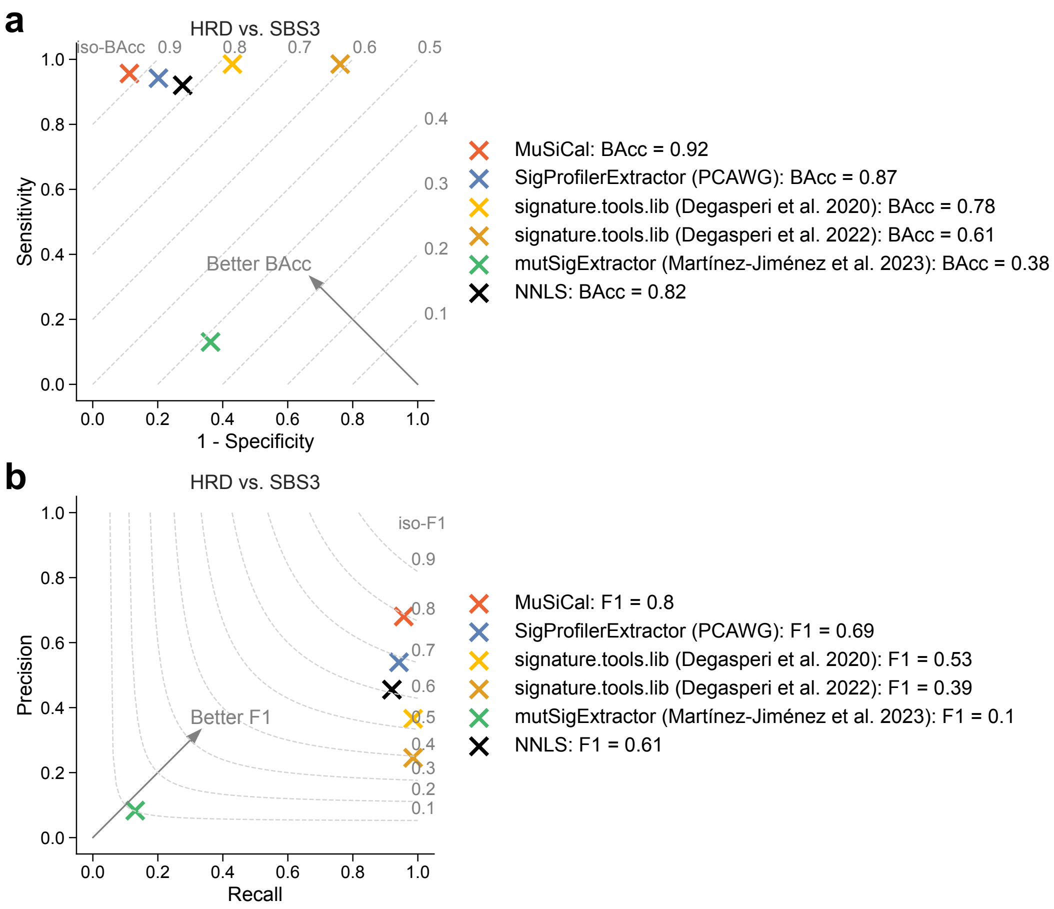

**Supplementary Fig. 14 MuSiCal-derived SBS3 assignment is more consistent with ground truth HRD status.** We compared SBS3 assignments from different methods to the ground truth HRD status in PCAWG breast, ovary, pancreas, and prostate tumors. The SBS3 assignments were binarized into SBS3 positive (nonzero) vs. negative (zero) labels before comparing to HRD labels. The HRDetect final classification from [13] was considered as ground truth. For SigProfilerExtractor, the final PCAWG signature assignment results from [4] were used. For signature.tools.lib, two versions of the results were used, one from [13], and the other from [14]. For NNLS, the spectrum of each sample was decomposed into the entire set of COSMIC signatures using NNLS. We also included results from [15]. **a.** Sensitivity vs. 1 - specificity. Dashed lines indicate iso-BAcc (balanced accuracy) lines. **b.** Precision vs. recall. Dashed curves indicate iso-F1 curves.

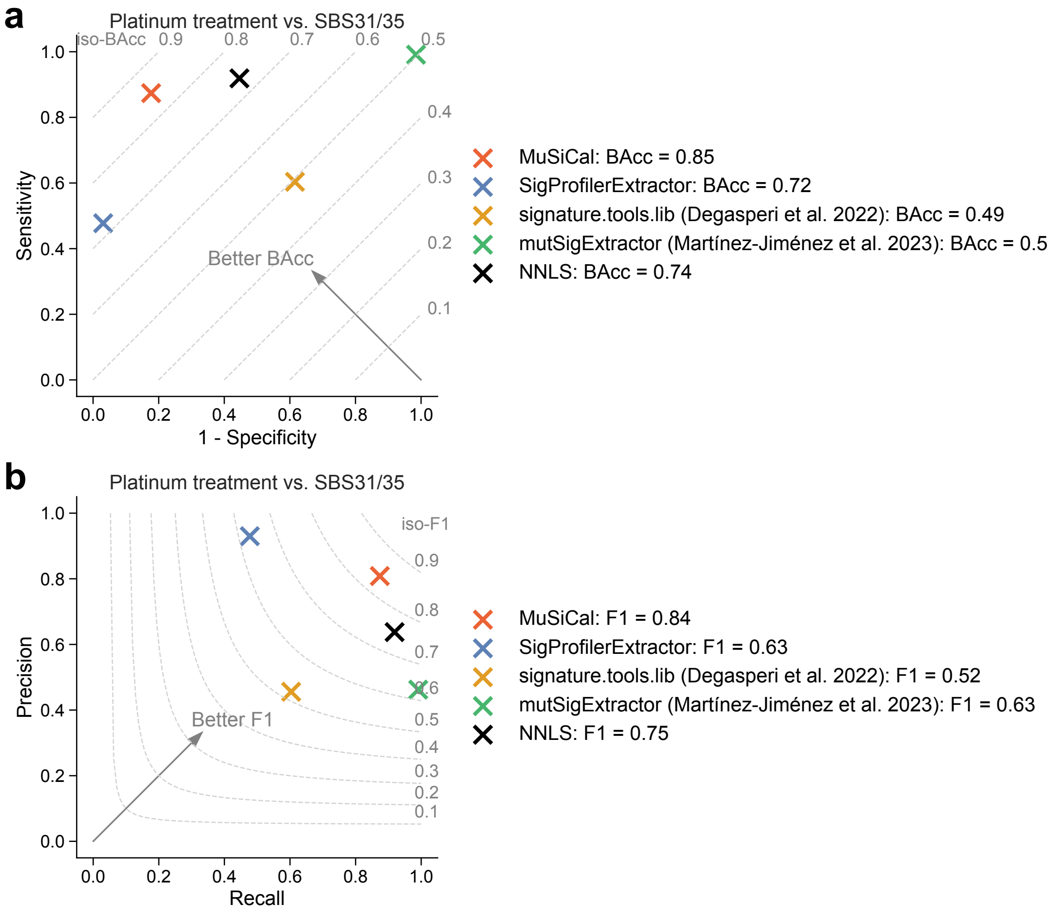

**Supplementary Fig. 15 MuSiCal-derived SBS31/35 assignment is more consistent with ground truth platinum treatment status.** We compared SBS31/35 assignments from different methods to the ground truth platinum treatment status in ovary tumors combined from Hartwig and PCAWG. n = 111 samples from Hartwig with before-biopsy treatments of Cisplatin, Oxaliplatin, or Carboplatin were considered platinum positive. n = 130 samples were considered platinum negative, which included n = 21 from Hartwig without before-biopsy treatments of Cisplatin, Oxaliplatin, or Carboplatin, and n = 109 from PCAWG (all assumed to be treatment naive). The SBS31/35 assignments were binarized into SBS31/35 positive (nonzero) vs. negative (zero) labels before comparing to platinum labels. MuSiCal and SigProfilerExtractor results were obtained by running the respective tool on this dataset. For signature.tools.lib, the signature assignment results from [14] were used. For NNLS, the spectrum of each sample was decomposed into the entire set of COSMIC signatures using NNLS. We also included results from the Hartwig paper itself [15] to be complete. **a.** Sensitivity vs. 1 - specificity. Dashed lines indicate iso-BAcc (balanced accuracy) lines. **b.** Precision vs. recall. Dashed curves indicate iso-F1 curves.

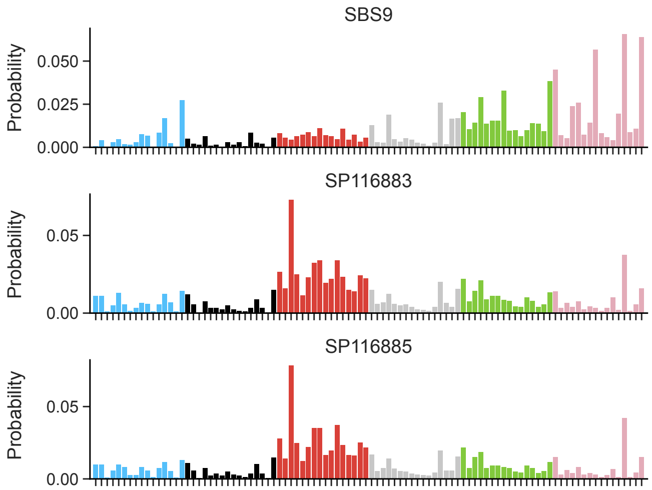

**Supplementary Fig. 16 Assignment of SBS9 in two Myeloid-MPN samples.** MuSiCal assigned SBS9 to 2 of the 56 Myeloid-MPN samples from PCAWG. The spectra of these two samples (SP116883 and SP116885) were shown together with SBS9. Both samples demonstrated characteristic peaks of SBS9 at T>A and T>G channels. However, SBS9 is known to be associated with polymerase eta activity during somatic hypermutation in lymphoid cells and is usually observed in post-germinal center lymphoid tumors. The assignment of SBS9 in these two myeloid tumors may thus represent a distinct mutational process that produces a similar signature in the trinucleotide context. Indeed, compared to SBS9, the T>G mutations in these two samples were specifically enriched at T[T>G]A.

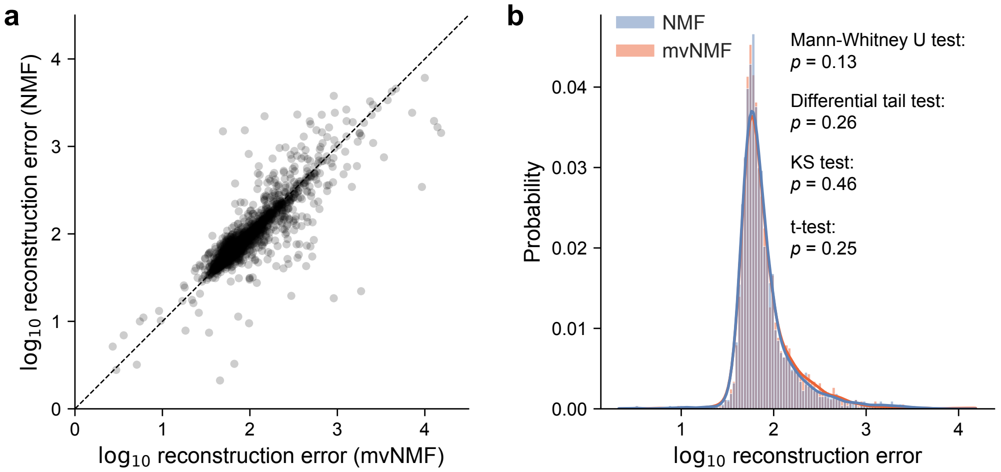

**Supplementary Fig. 17 Comparison of reconstruction errors of NMF and mvNMF solutions on PCAWG data. a.** Per-sample reconstruction errors of NMF vs. mvNMF solutions. Most of the data points were close to the diagonal (dashed line), suggesting that NMF and mvNMF resulted in similar reconstruction errors (Pearson $r = 0.88$, $p < 10^{-300}$). Slightly more than half (57%) of the samples had larger reconstruction errors from mvNMF. To ensure a fair comparison, the NMF solution was forced to have the same number of signatures as the mvNMF solution for the corresponding tumor type. **b.** Distribution of the per-sample reconstruction errors for NMF and mvNMF solutions separately. $P$-values from four different statistical tests (all two-sided) were annotated for testing the difference between the two distributions. None of them were statistically significant.

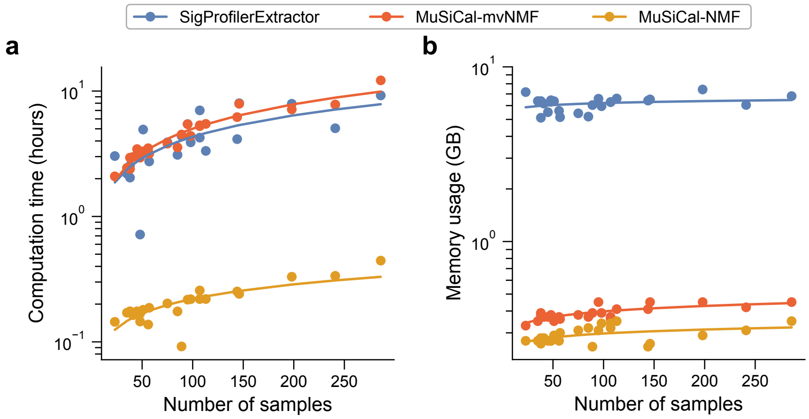

**Supplementary Fig. 18 Computational cost of MuSiCal in comparison to SigProfilerExtractor.** MuSiCal (in both mvNMF and NMF modes) and SigProfilerExtractor (based on NMF) were run on each PCAWG tumor type separately for *de novo* signature discovery, and the corresponding computation time and memory usage were shown. MuSiCal was run with 10 CPUs on a high-performance cluster, and SigProfilerExtractor was run with 12 CPUs. **a.** Computation time (in hours) for *de novo* signature discovery was plotted against the number of samples. Each dot represents a PCAWG tumor type. Solid lines represent linear fits in the log space, i.e., $\log(t) \sim \log(n)$, where $t$ denotes computation time, and $n$ number of samples. MuSiCal with mvNMF and SigProfilerExtractor were comparable in computation time for PCAWG tumor types, although MuSiCal with mvNMF scaled slightly worse ($t \propto n^{0.65}$ for MuSiCal with mvNMF and $t \propto n^{0.57}$ for SigProfilerExtractor). MuSiCal with NMF was much faster and scaled the best ($t \propto n^{0.39}$). **b.** Same as panel (a) but for memory usage (in GB). MuSiCal with either mvNMF or NMF required considerably less memory than SigProfilerExtractor.

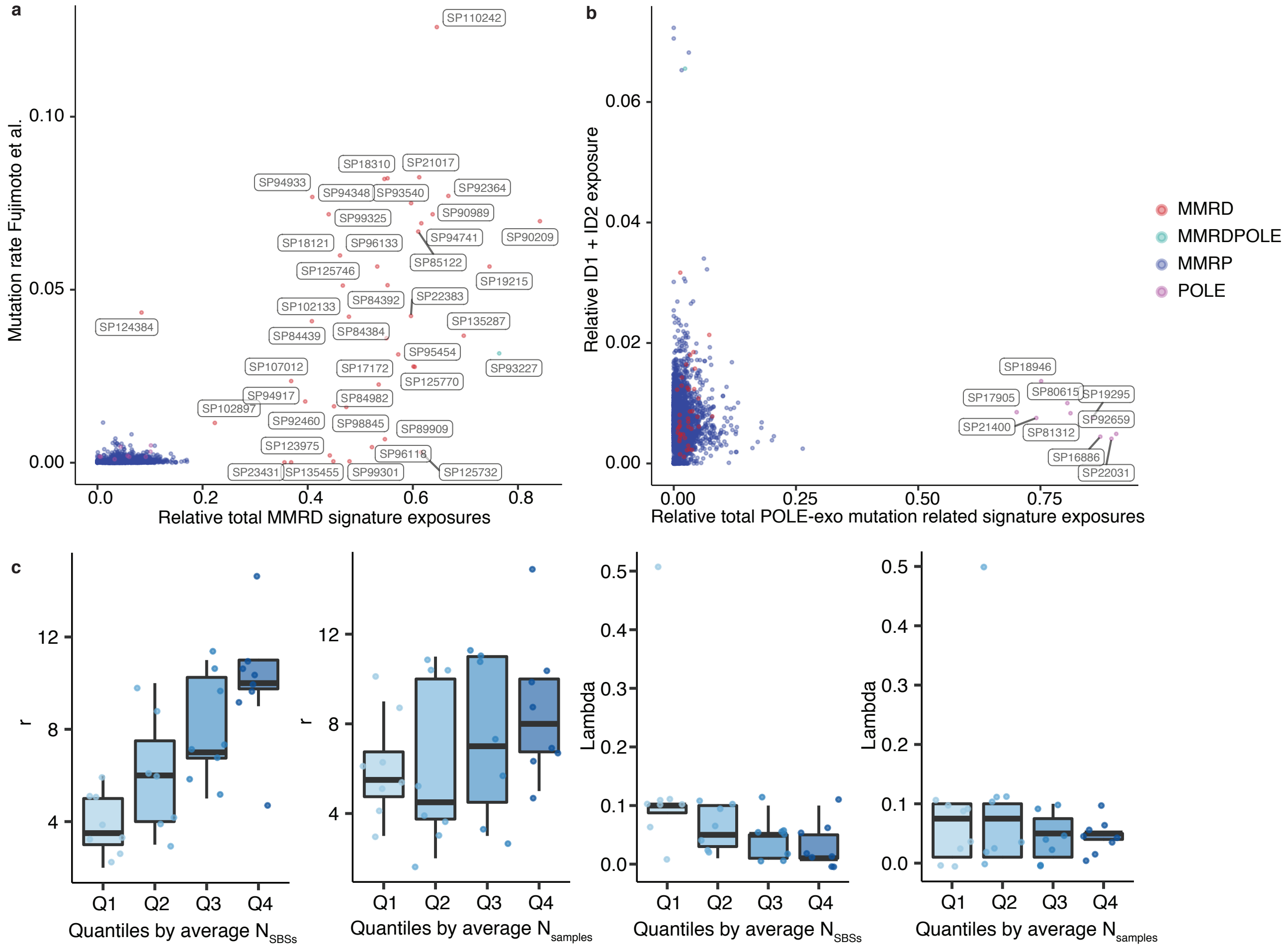

**Supplementary Fig. 19 Details of PCAWG reanalysis for SBS signatures. a.** Microsatellite mutation rate (as obtained from [16]) vs. relative exposure of MMRD-related SBS signatures (SBS6, 14, 15, 20, 21, 26, 44) for PCAWG samples. A clear separation is observed between two classes of samples, which we identify as MMRD and MMRP, respectively. SBS14 exposure alone is used to define the single sample with concurrent MMRD and POLE-exo mutations. **b.** Relative exposure of ID1 + ID2 vs. relative exposure of SBS signatures associated with POLE-exo mutations (SBS10a-d and SBS28). A clear separation is observed along the x-axis, which is used to identify tumors with POLE-exo mutations. **c.** Dependence of the selected number of signatures $r$ and the optimal mvNMF regularization parameter $\tilde{\lambda}$ on the number of samples and the average number of SBSs.
